# Reference materials for SARS-CoV-2 molecular diagnostic quality control: validation of encapsulated synthetic RNAs for room temperature storage and shipping

**DOI:** 10.1101/2023.08.28.555008

**Authors:** Marthe Colotte, Aurélie Luis, Delphine Coudy, Sophie Tuffet, Isabelle Robene, Babbitha Fenelon, Emmanuel Jouen, Nicolas Leveque, Luc Deroche, Sophie Alain, Dorian Plumelle, Camille Tumiotto, Laurent Busson, Marie-Edith Lafon, Jacques Bonnet

## Abstract

The Coronavirus pandemic unveiled the unprecedented need for diagnostic tests to rapidly detect the presence of pathogens in the population. Real-time RT-PCR and other nucleic acid amplification techniques are accurate and sensitive molecular techniques that necessitate positive controls. To meet this need, Twist Bioscience has developed and released synthetic RNA controls. However, RNA is an inherently unstable molecule needing cold storage, costly shipping, and resource-intensive logistics. Imagene provides a solution to this problem by encapsulating dehydrated RNA inside metallic capsules filled with anhydrous argon, allowing room temperature and eco-friendly storage and shipping. Here, RNA controls produced by Twist were encapsulated (RNAshells) and distributed to several laboratories that used them for COVID-19 detection tests by amplification. One RT-LAMP procedure, four different RT-PCR devices and 6 different PCR kits were used. The amplification targets were genes E, N; RdRp, Sarbeco-E and Orf1a/b. RNA retrieval was satisfactory, and the detection was reproducible. RNA stability was checked by accelerated aging. The results for a 10-year equivalent storage time at 25 °C were not significantly different from those for unaged samples. This room temperature RNA stability allows the preparation and distribution of large strategic batches which can be stored for a long time and used for standardization processes between detection sites. Moreover, it makes it also possible to use these controls for single use and in the field where large temperature differences can occur. Consequently, this type of encapsulated RNA controls, processed at room temperature, can be used as reference materials for the SARS-Cov-2 virus as well as for other pathogens detection.

## Introduction

The Coronavirus pandemic has led to an unprecedented need for diagnostic tests to rapidly detect the presence of this pathogen in the population. Real-time RT-PCR and RT-LAMP amplification techniques proved to be accurate and sensitive molecular tools for virological diagnosis [1,2,3,4,5].

A key aspect of any ongoing pathogen detection effort is using quality control measures for a wide range of applications from diagnostic assay development to day-to-day testing. This includes verifying and validating diagnostic tests of both Next-Generation Sequencing (NGS) and reverse transcription-amplification (RT-PCR or RT-LAMP) assays [6,7,8,9]. Controls based on viral nucleic acids extracted from infected patients or live viruses propagated in cell culture have safety and security concerns. Synthetic RNA controls, like those generated through gene synthesis by Twist Bioscience, mitigate the biohazard risk and broaden access across diverse strains [10]. Additionally, as DNA synthesis is a readily available technology, new RNA controls can be rapidly developed in response to the emergence of new strains or pathogens.

However, RNA is an inherently unstable molecule. In solution or dry i), RNA rapidly degrades through hydrolysis or oxidation (for a review, see [11]). This instability presents storage and cost challenges for laboratories that require large stocks of control RNA (for a review see [2]). Imagene’s technology consists of confining dehydrated RNA, in the presence of proprietary stabilizing solutions, inside metallic capsules filled with anhydrous argon (RNAshells®), keeping away the samples from deleterious factors. Thus, it offers a solution to the stability problem by allowing extensive and eco-friendly storage and shipping a room temperature [2].

Here, Imagene encapsulated synthetic RNA controls (RNAshells) produced by Twist Bioscience. These controls were then distributed to five laboratories for fitness-for-purpose analysis. The objective was to verify, first, that the detection of the virus was reliable and second that the RNA stability was insured for a minimum of a 10-year storage period at room temperature. For that, we used four different RT-PCR devices with six different PCR kits and one LAMP procedure on RNA from three SARS-Cov-2 variants and five gene targets.

## Material and methods

### Workflow

The RNA samples synthetized by Twist Bioscience were sent on dry ice to Imagene. One part of these samples was aliquoted and sent on dry ice to Bordeaux CHU. The other part was aliquoted and encapsulated. A set of these aliquots was heated to simulate 10 years of room temperature storage and sent to Bordeaux CHU. The other aliquots were sent to Bordeaux, Limoges, Poitiers CHUs, the CIRAD Réunion and a private clinical laboratory (Plumelle, Salon de Provence).

The experimental strategy we followed is summarized in Fig 1.

**Fig 1.**
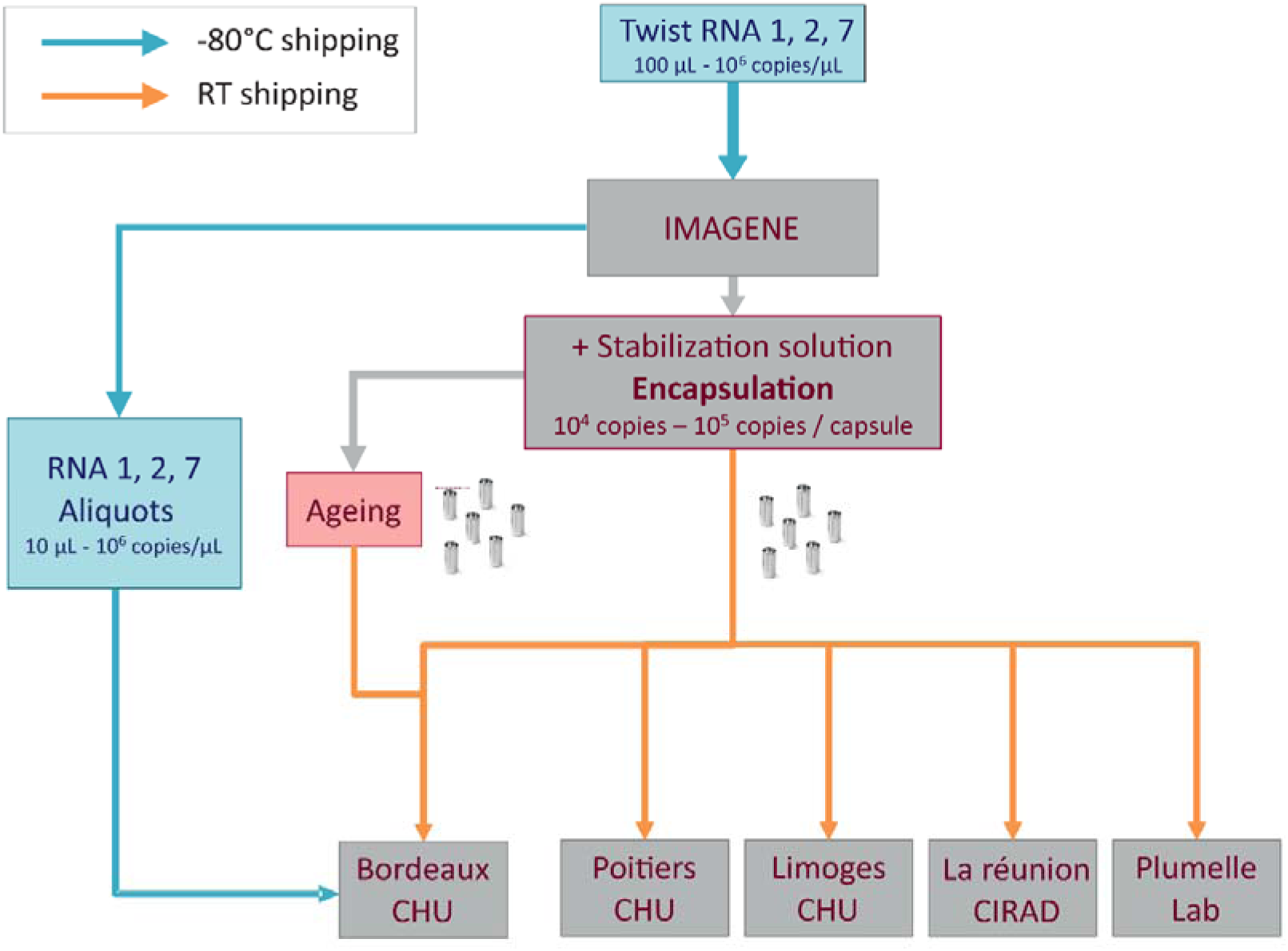
Experimental strategy for the preparation, distribution and validation of RNAshell® encapsulated RNA controls for SARS-CoV-2 virus detection.

### RNA Twist RNA controls

Synthetic SARS-CoV-2 RNA control molecules for three SARS-CoV-2 variants were graciously supplied by Twist Bioscience. Details are given Table 1.

**Table 1.**
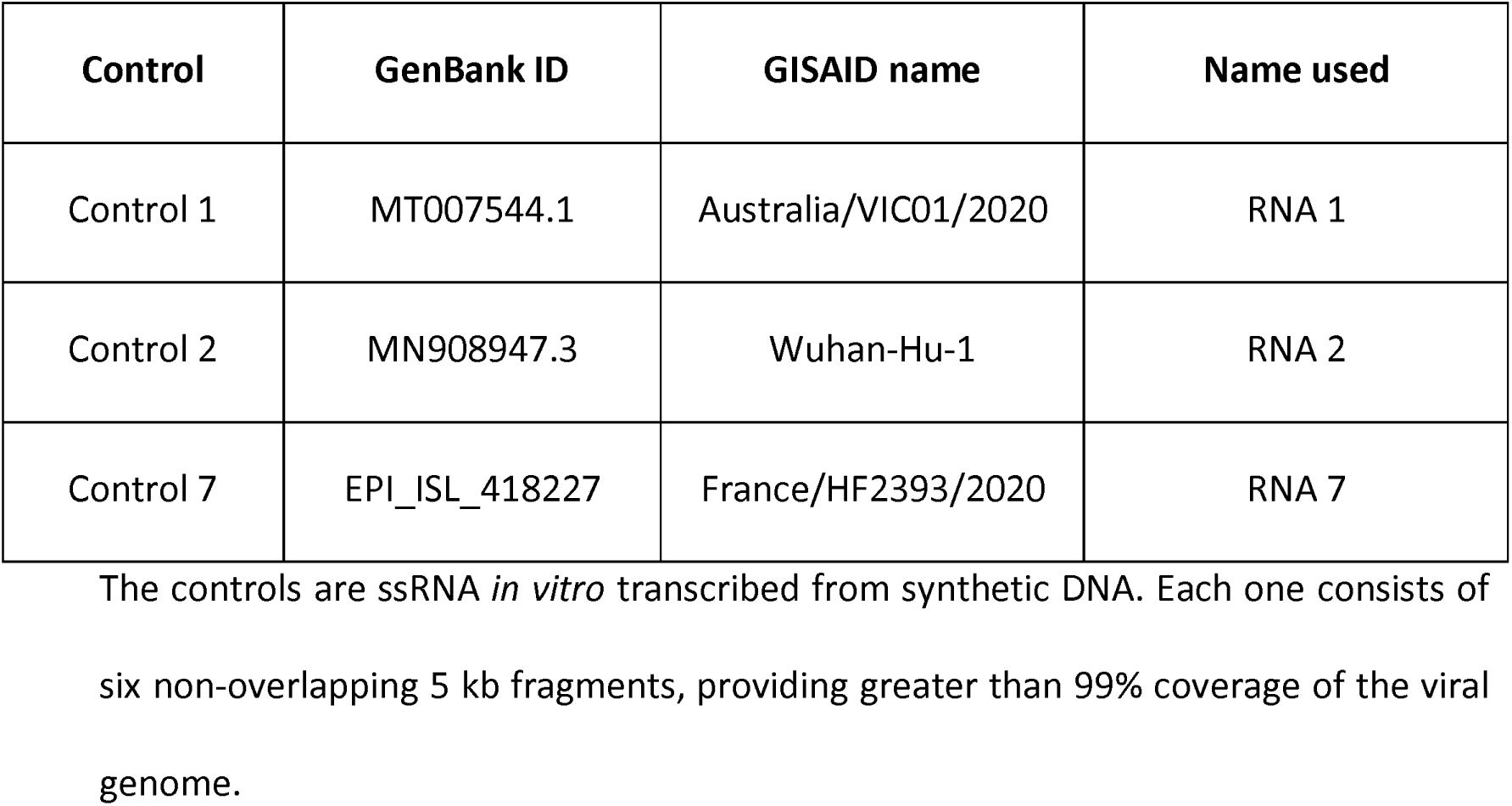
Synthetic SARS-CoV-2 RNA controls used in this study.

### RNA encapsulation and accelerated ageing

RNA controls were sent to Imagene over dry ice as 100 μL solutions at a concentration of approximately 10^6^ copies per microliter [10]. They were distributed in 10 µL aliquots and stored at –80°C. Aliquots were thawed, diluted with an Imagene proprietary stabilization solution and distributed into glass inserts fitted in capsules to obtain either 10^4^ copies (low) or 10^5^ copies (high) per capsule.

RNA samples were vacuum-dried and encapsulated under anhydrous and anoxic argon/helium according to the process developed by Imagene [12,13] to give RNAshells®. A set of 3 capsules of each variant RNA and each copy number (a total of 18 capsules) were heated at 90 °C for 16 h to simulate 10 years of storage at 25 °C according to our previously published Arrhenius equation for encapsulated RNA [11]. RNAshells® were then shipped at room temperature to the participating laboratories.

### RNAshell® opening and RNA rehydration

The RNAshells® were opened with a portable ShellOpener^TM^ supplied by Imagene following the given instructions. After opening, a given volume of RNAse-free ultrapure water was added in the capsule and the RNA was left 10-20 minutes on the bench to rehydrate. At that point, the RNA was ready for analysis (RT-qPCR or RT-LAMP). The different volumes used to rehydrate and analyze the RNA by RT-qPCR in the different laboratories are given in Table 2.

**Table 2.**
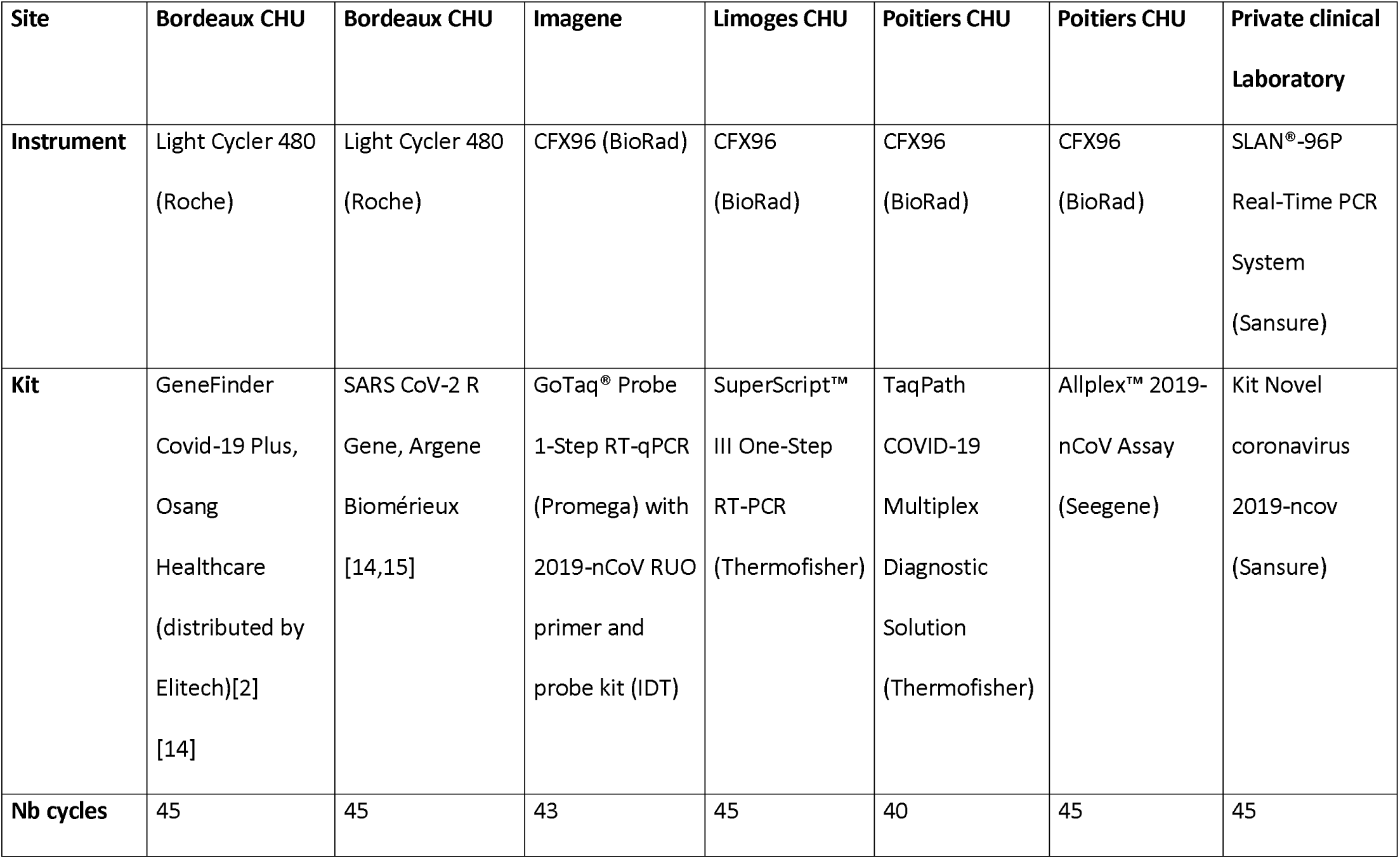
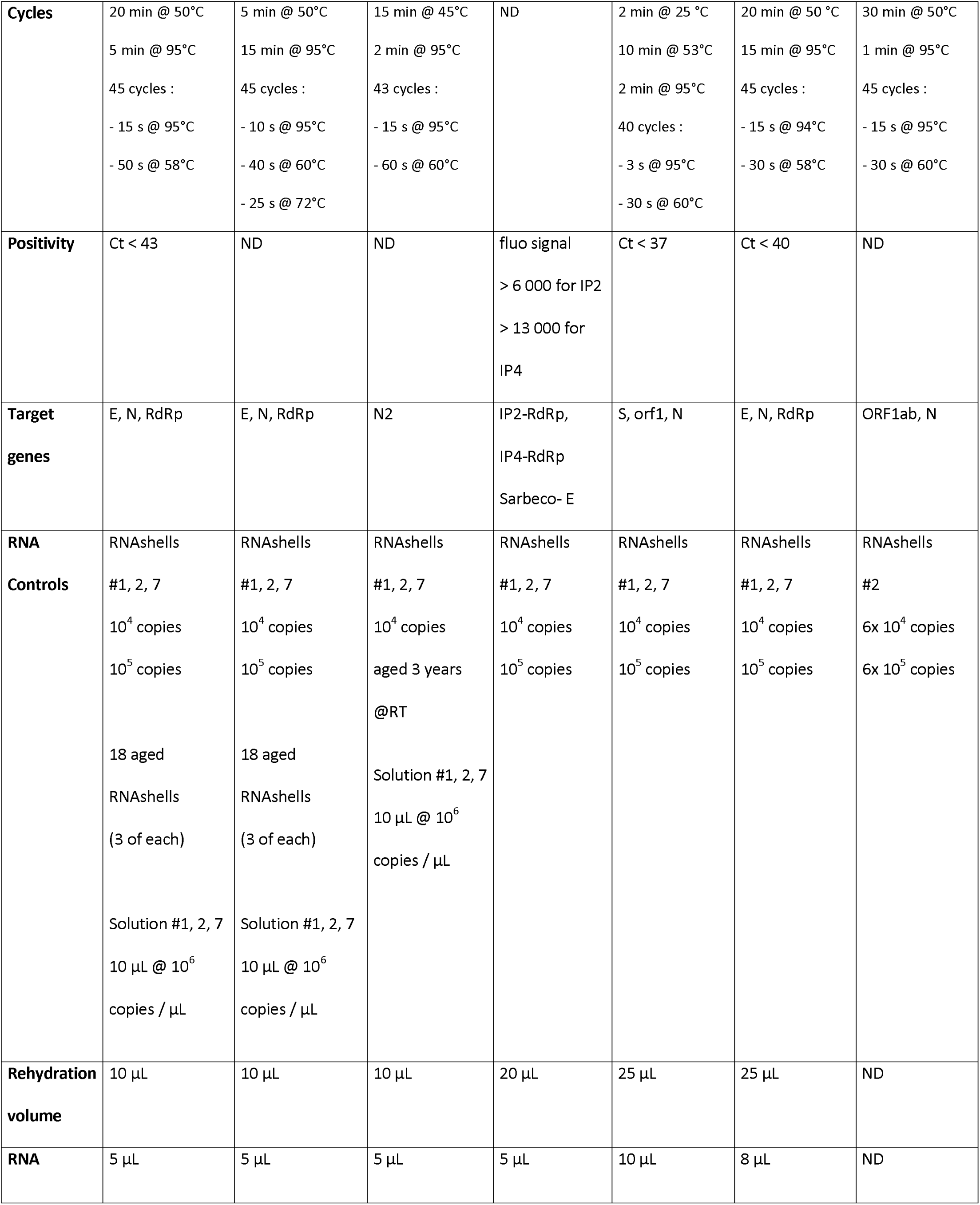
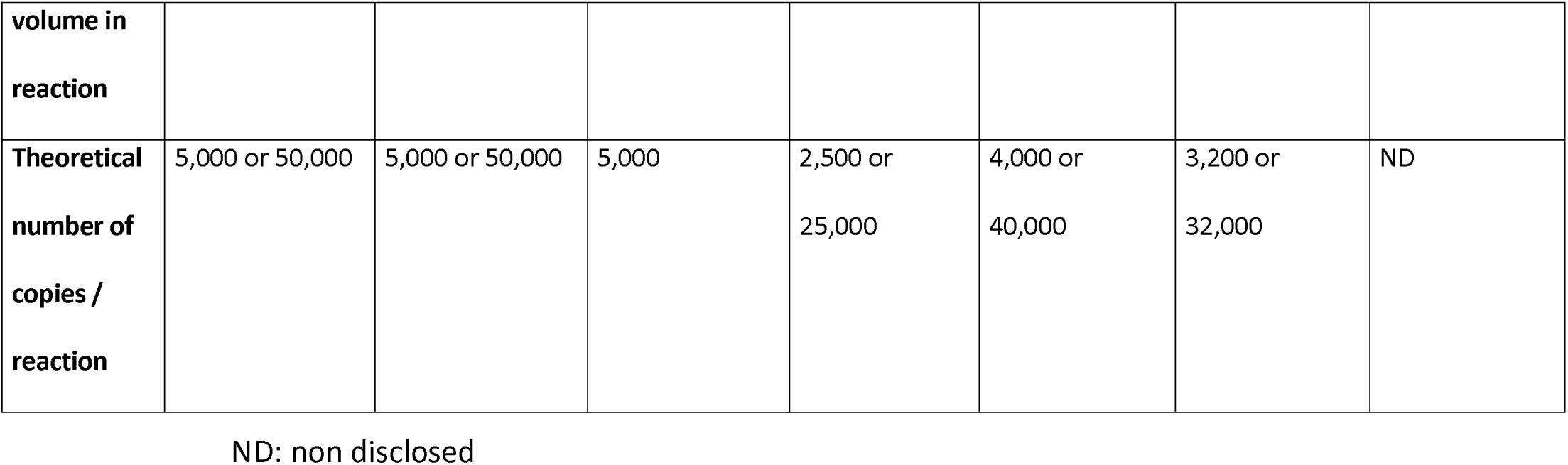
Summary of the material and methods for the different RT-qPCR analysis.

### Analysis of RNA samples in solution

The non-diluted and non-encapsulated aliquots of RNA 1, 2 and 7, at approximately 10^6^ copies per µL, were prepared and sent by Imagene on dry ice to Bordeaux CHU. The samples were thawed and diluted to match the concentrations of the rehydrated capsules (here, 10^3^ and 10^4^ copies per µL). They were analyzed in parallel by RT-qPCR (see Table 2).

### Control for DNA contamination

The “No-RT” controls were performed to look for DNA contaminations in the provided RNA controls. They consisted in preparing the RT-qPCR mix with all reagents and RNAs but without the RT enzyme. They have been run on 3-year-old capsules and –80°C frozen controls for RNA 1, 2 and 7 (see Table 2, Imagene).

### RT-qPCR analysis

The different available RT-qPCR materials and methods used in the study are summed up in Table 2.

### RT-LAMP analysis

Here, the RT-LAMP system was first tested on a dilution series of control RNA 2 Low or High, rehydrated with 50 µL of RNAse-free pure water. For Control 2 Low: a single analysis was done for 1000 copies/reaction, 2 replicates were done for 500 and 250 copies/reaction and 3 replicates for 100, 50 and 25 copies/reaction. For Control 2 High: 3 replicates were done for 1000 copies/reaction and 4 replicates for 200, 40 and 8 copies/reaction.

To test the encapsulated RNA 2 as a positive control in the field, 60 independent RT-LAMP runs were performed for the presence of SARS-CoV-2. Each run included 14 samples, a negative control and a positive control. The positive control consisted of RNA 2 rehydrated with water and further diluted in *Amies* medium to have a final concentration of 1000 copies/reaction.

RT-LAMP was performed according to the protocol RunCov developed by the CIRAD-UMR PVBMT (3P) and validated by the National Reference Center [16]. The reactions were performed in a 25 μL total reaction volume containing 15 μL of ISO-DR004-RT300 Isothermal Mastermix (OptiGene Ltd, Horsham, United Kingdom), 2.475 μL of each primer pair (N and S), 0.05 μL (0.5 units) of AMV transcriptase (Promega, Charbonnières-les-Bains, France) and 5 μL of template RNA. LAMP reactions were run on a Genie II instrument (OptiGene) at 65°C for 25 min followed by an annealing step during which the temperature was decreased from 92°C to 78°C at a speed of 0.05°C/s to determine the annealing temperature (Ta) of the produced amplicon.

### Statistical analysis

All statistical analyses including ANOVA, Fisher Test and previous verification of data normality (histogram and normal: Q-Q plot), were performed using the R statistical software, version 4.1.1 (2021-08-10; R Development Core Team, Vienna, Austria) with the different packages: stats, multcomp, emmeans and ggplot2).

## Results and discussion

### Objectives and strategy

The aim of this work was to validate whether encapsulated RNA controls stored and shipped at room temperature were fit for purpose as reference materials for internal quality control of SARS-CoV-2 detection.

Bordeaux CHU received non-diluted, non-encapsulated, frozen samples in solution, to compare the controls before and after encapsulation (i.e. to check the potential effect of the process and stabilization solution). They also received the aged RNAshells® to validate that 10 years of storage at room temperature did not alter the performance of the encapsulated controls. Two other laboratories (Poitiers CHU and Limoges CHU) received RNAshells® to verify that the controls gave the same performances when run with other RT-qPCR techniques. The CIRAD-UMR PVBMT (3P) laboratory tested the controls in multiple experiments while developing a RT-LAMP test for field use. Finally, RNAshells® were sent to a private clinical laboratory to check that the capsules can be routinely used by any operator.

First we ran controls, on the one hand for detecting eventual DNA contamination in RNA samples and on the other hand to check the performance of non encapsulated controls.

### Controls for DNA contamination

We ran RT-qPCR minus reverse transcriptase on RNA 1, 2, 7 since these RNA were prepared by reverse transcription from DNA templates and DNA is much more stable than RNA, even though a DNA contamination large enough to bias the RT-PCR results is rather unlikely considering the way RNA samples have been synthetized. Indeed, in a typical in vitro transcription synthesis system (for instance [17]) 1) there are at least 20 RNA copies generated from one DNA template molecule, 2) after synthesis, there is a DNase treatment and 3) there is a final purification step.

The results are given Fig 2.

**Fig 2.**
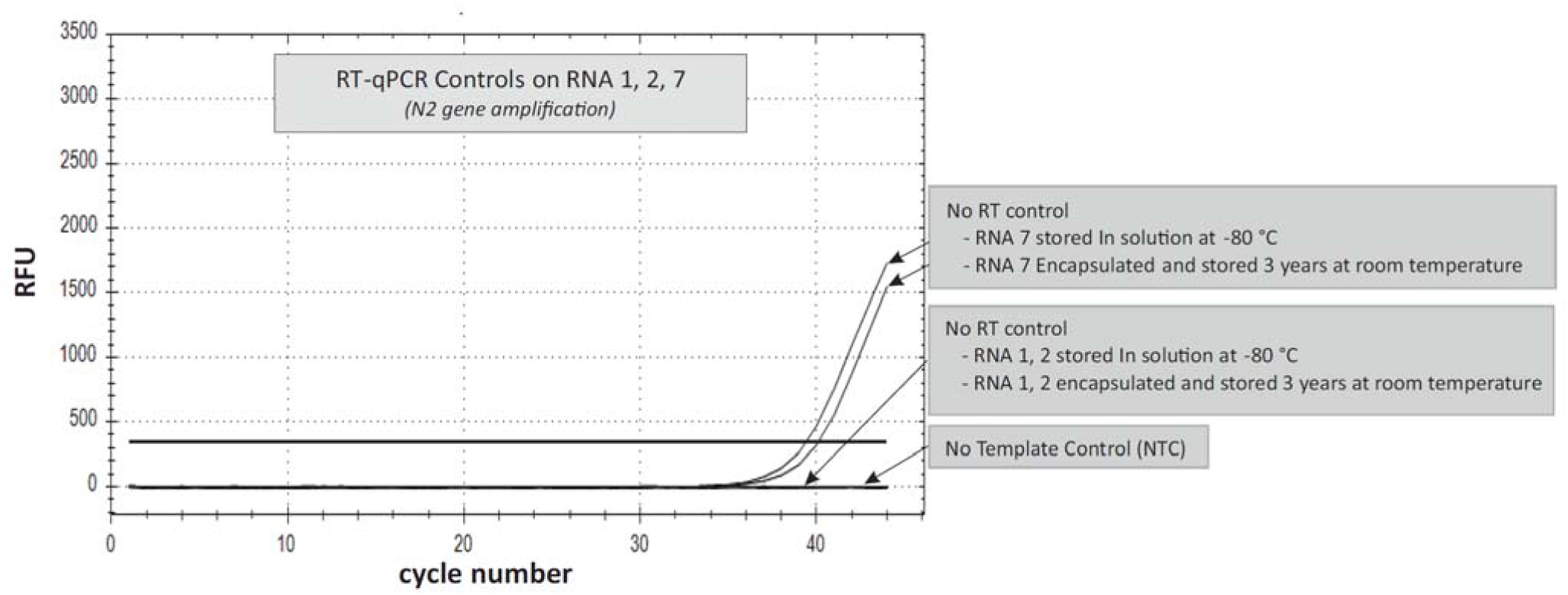
RT-qPCR controls for DNA contamination. The amplifications, with or without reverse transcription, were conducted as described in materials and methods.

The no RT controls for RNA 1 and RNA 2 were negative but RNA 7 actually showed the presence of a slight DNA contamination. However, the measured 39-40 Ct was clearly beyond the 25-30 Ct of all (excepted 2, discussed Fig 7) Ct values measured in this study and could not have any impact.

### Controls in solution

The results of these experiments are presented Fig 3.

**Fig 3.**
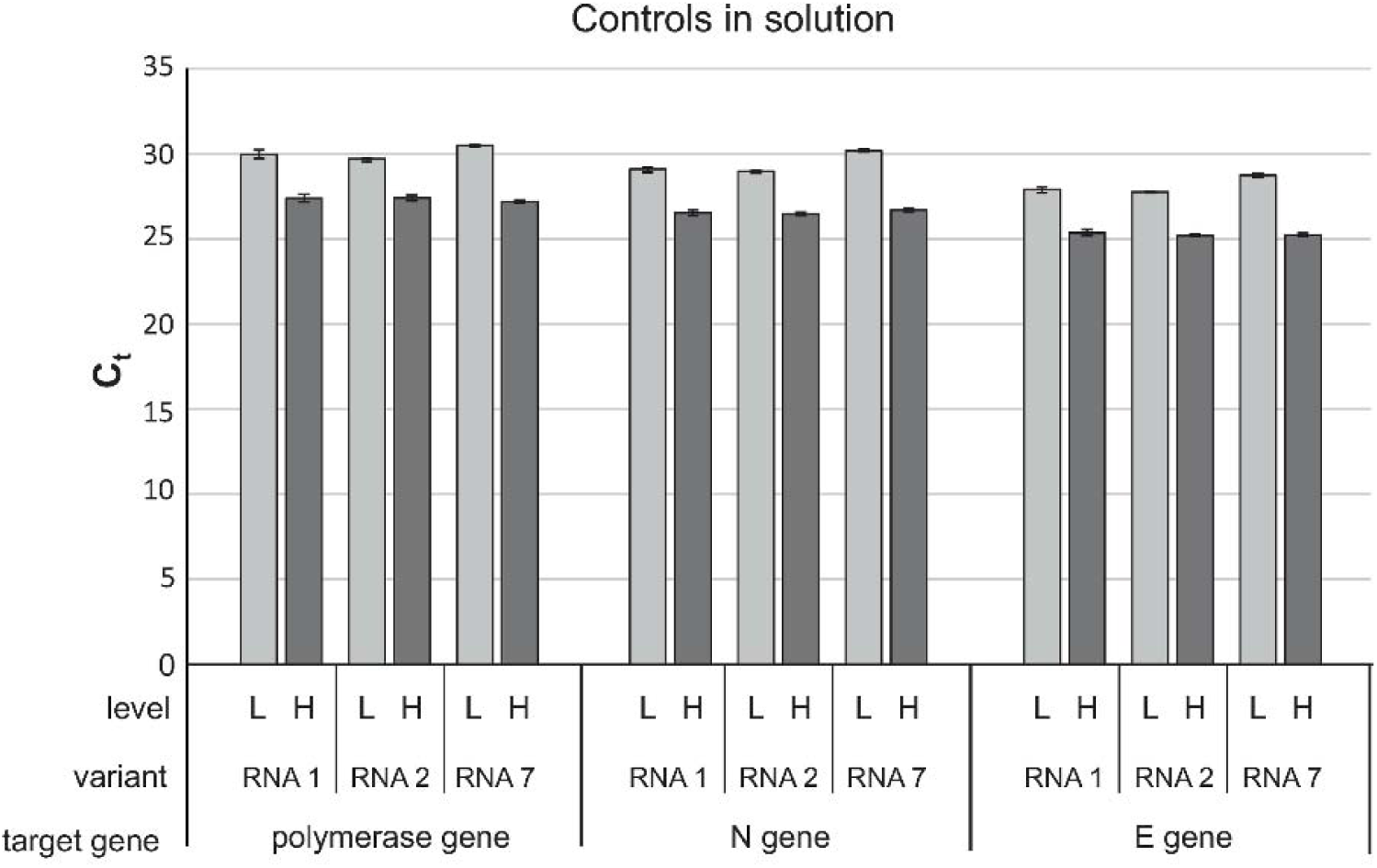
RT-qPCR for controls in solution. . The controls were thawed and diluted at reception. Measures were done at Bordeaux CHU on 3 variants RNA (RNA 1, 2, 7), 2 copy numbers: 5,000 (low, L) and 50,000 (high, H) copies/reaction, 3 target genes (polymerase gene, N gene and E gene).

All the controls gave positive results. For a given low or high concentration and a given target gene, the Ct were comparable between the different variants (p = 0.647). The highest Ct differences came from the gene targets (p=7.79 x 10^-4^), probably due to differences in PCR efficiency.

The chosen copy numbers gave Ct values in a suitable range (around 30 and 27 respectively for the low and high controls), clearly different from the negative control (above 40-45). The Ct were also not too low. This is advantageous because a positive control with too many more copies than in patient samples could result in carry-over contamination in the nearby wells of the PCR plate leading to false positive samples.

### RNA controls encapsulation and accelerated aging

Then, we needed to verify, that encapsulation and retrieval did not induce alterations in the controls and that these controls could be stored at room temperature for a long time in RNAshell®, so a set of capsules were submitted to an accelerated aging by heating for a period simulating a storage period of 10 years at 25°C according to our previously published Arrhenius study [11]. The results are shown Fig 4.

**Fig. 4.**
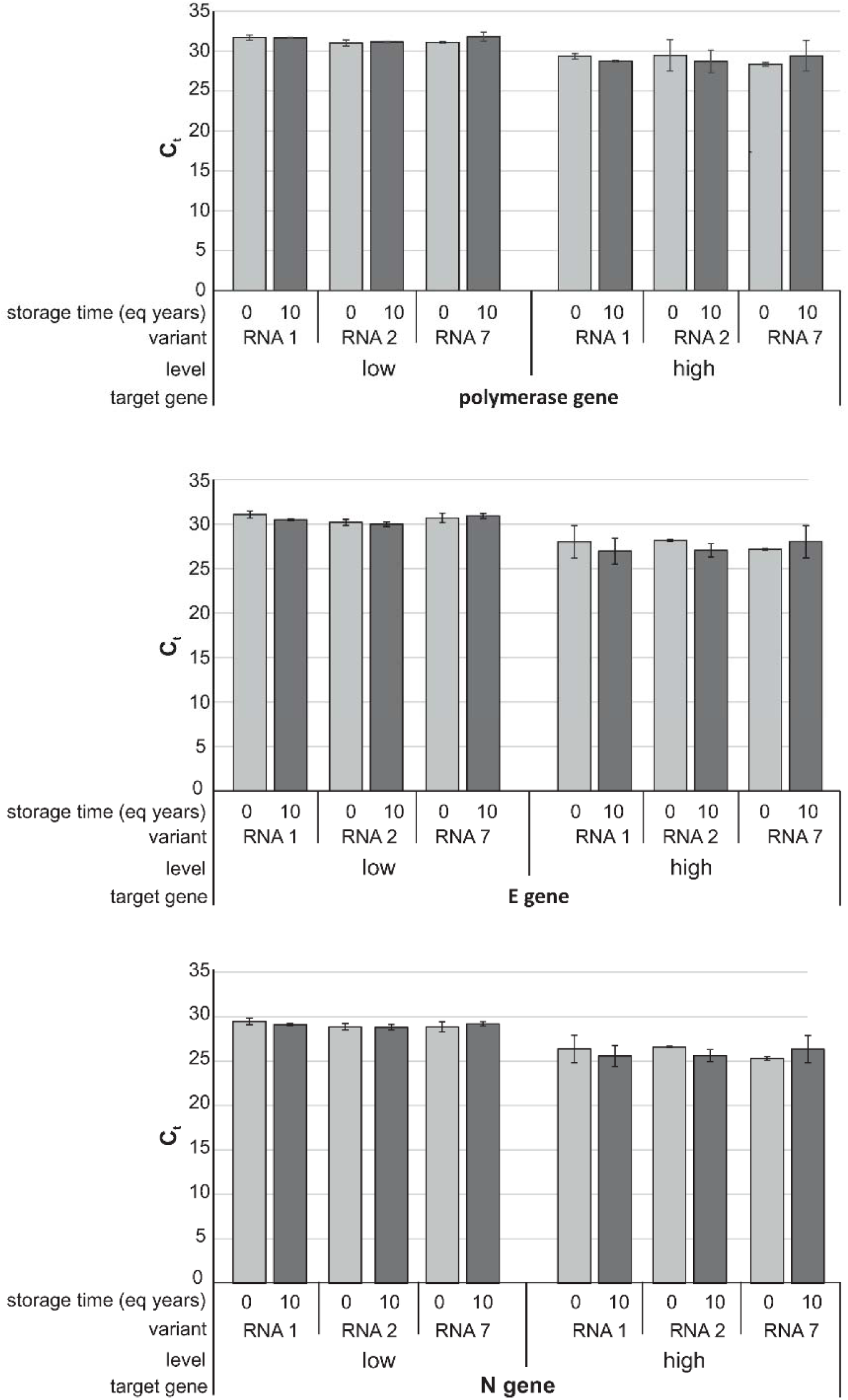
RT-qPCR on RNAshell®-encapsulated and aged controls. RT-qPCR measures were done by Bordeaux CHU on 3 variants RNA (RNA 1, 2, 7), 2 copy numbers: 5,000 (low, L) and 50,000 (high, H) copies/reaction, 3 target genes (polymerase gene, N gene and E gene) and after 2 storage times: time 0 post-encapsulation and after heating for a period equivalent to 10 years at room temperature (0 and 10).

All analyzed RNA, whatever the variant, target gene or storage time were successfully amplified.

To analyze the extent and sources of variations on these experiments we used variant 1 for a group analysis. The results are shown Fig 5.

**Fig 5.**
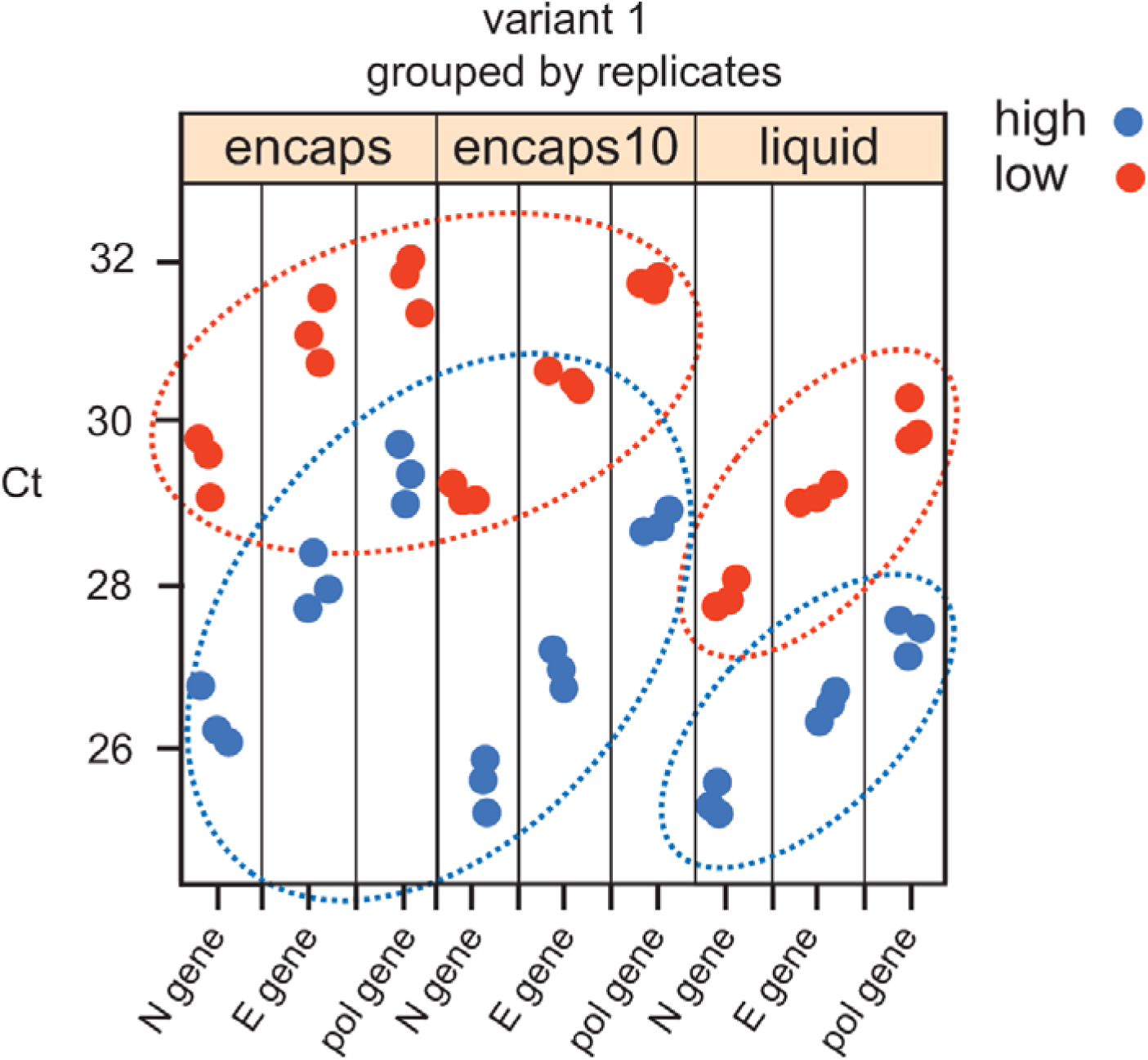
Group analysis for Ct on variant 1 controls. Ct (3 replicates) are shown according to: a) the state: encapsulated (encaps), encapsulated and aged (encaps 10) or in solution (liquid) and b) the number of copies (5,000 (low) and 50,000 (high) of each target gene. Groups of Ct values from the same state (dry vs liquid) and copy number are circled.

The encapsulated controls and the controls in solution show significant Ct differences (p=0.001999), due partly to the encapsulation and retrieval steps, including the addition of a stabilization solution, but more probably to the fact that the controls in solution did not come from the same diluted samples. However, the Ct values of encapsulated RNA are maintained in the same suitable range, meaning that the encapsulation process did not lead to RNA loss, degradation or PCR inhibition to an extent that would make RNAshell®-encapsulated RNA unfit for purpose.

On the other side, no significant difference was observed between 0– and 10-years RNAshells® (Fig 3, p=0.7606) meaning that the simulated long-term storage of RNA did not lead to significant degradation, as expected by the complete Arrhenius study we previously performed [12]. One can therefore consider that the encapsulated RNA controls are stable over at least 10 years at room temperature and thus remain suitable reference materials.

To go further, we ran a real time stability measurement on encapsulated controls over 3 years at room temperature. The results, compared to RNA stored frozen at –80 °C are shown on Fig 6.

**Fig 6.**
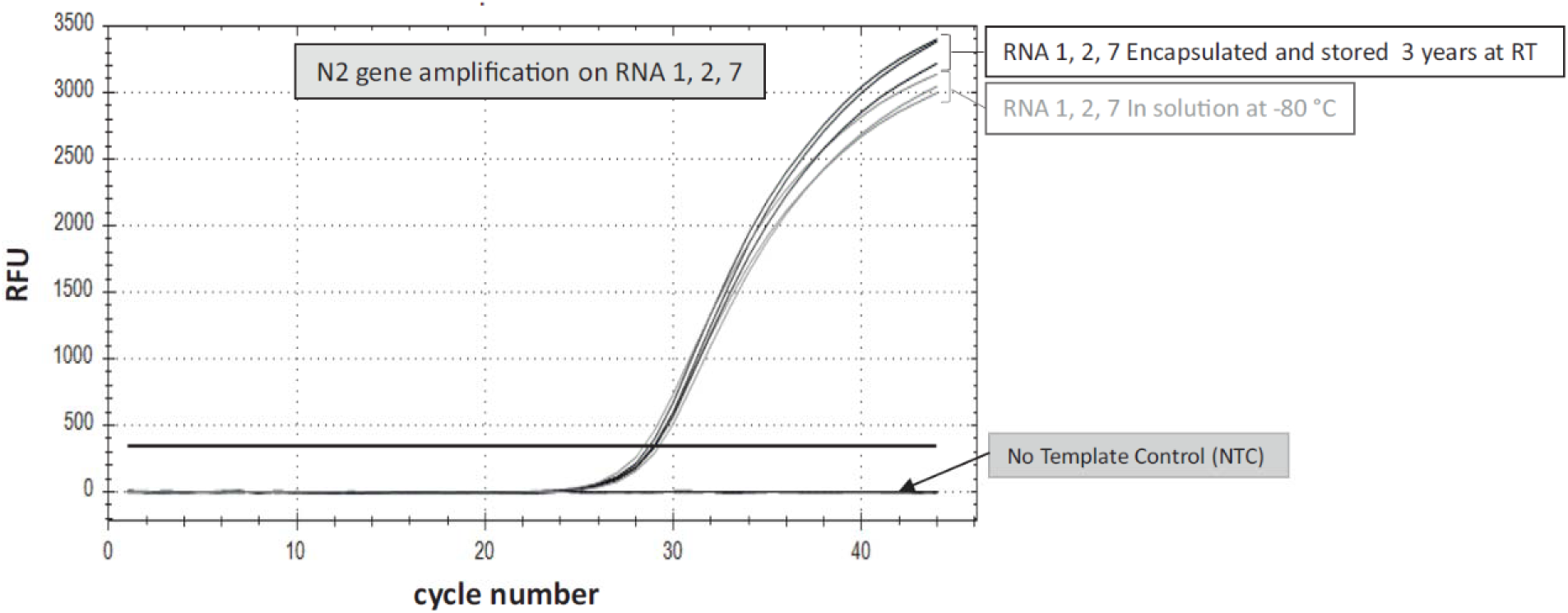
Room temperature stability of encapsulated RNA. These measurements were done on RNA 1, 2, 7. The RT-qPCR were done according to material and methods.

It can be seen that there is no Ct difference between the RNA kept 3 years frozen in solution at –80 °C and the encapsulated RNA stored at room temperature for the same time. The Ct are comparable to the Ct obtained after encapsulation (Fig. 4).

### Compatibility with other RT-qPCR kits and instruments

To broaden the validation, capsules were distributed to laboratories that performed tests with three other RT-amplification techniques, including the method used by the National Reference Center for SARS-CoV 2 (on RdRp IP2 and IP4 regions). The results are shown Fig 7.

**Fig 7.**
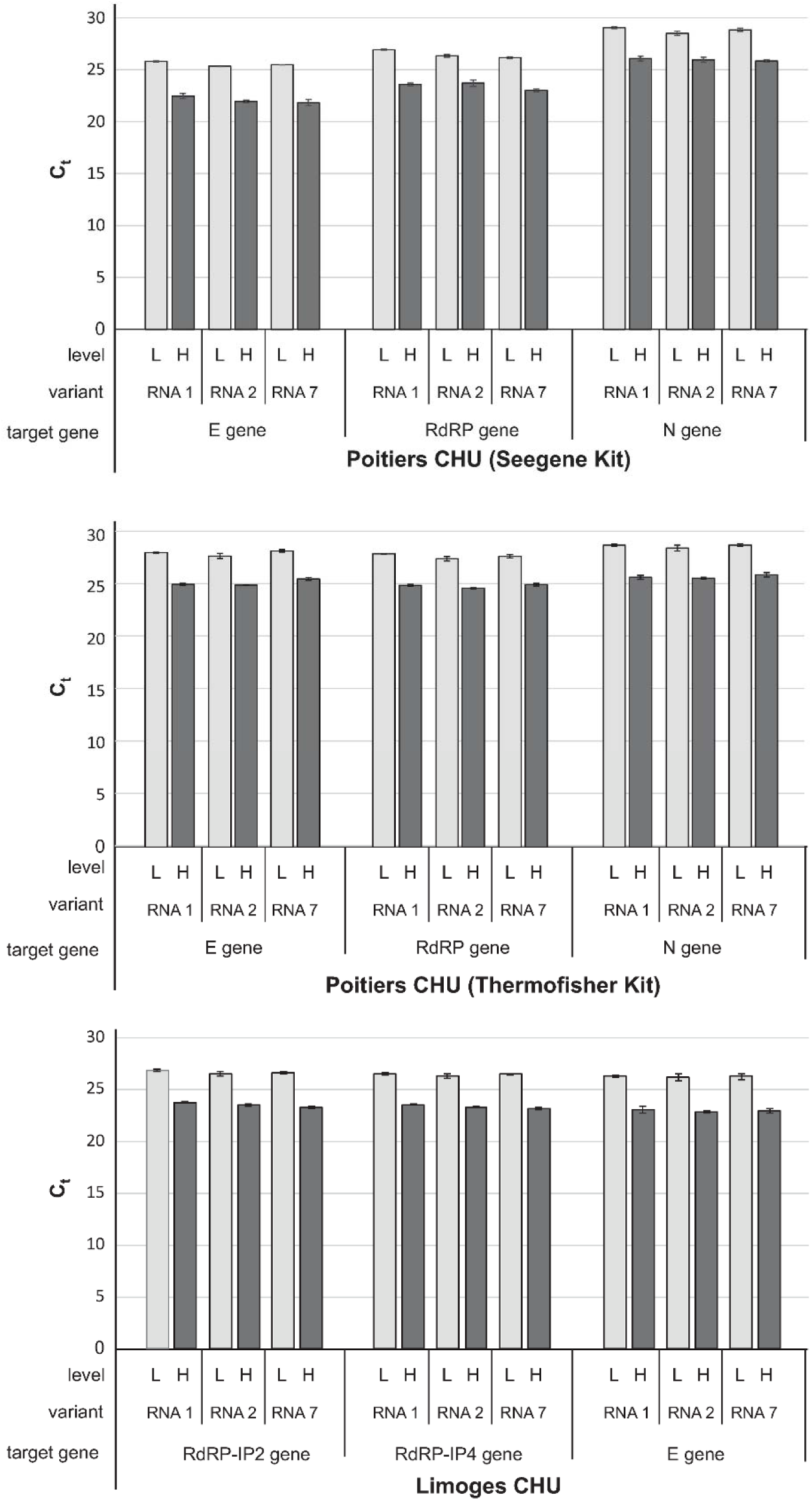
RT-qPCR results from Limoges and Poitiers CHU. For Limoges CHU, the RT-qPCR measures were done on 3 variants RNA (RNA 1, 2 and 7), 2 copy numbers: 2,500 (Low, L) and 25,000 (High, H) copies/reaction), and 3 target genes (RdRp-IP2, RdRp-IP4 and E genes). For Poitiers CHU, RNAshells® were rehydrated with 25 µL of water. Eight µL were used for the Seegene kit and 10 µL for the Thermofisher kit. Ct values were obtained by RT-qPCR measures done on 3 variants RNA (RNA 1, 2 and 7), 3 target genes (E, N and RdRp genes), with 2 copy numbers: 10,000 (Low, L) and 100,000 copies/capsule (high, H) giving 3,200 and 32,000 copies per reaction for the Seegene kit and 4,000 and 40,000 copies per reaction for the Thermofisher kit. For each technique, the control capsules were run in 3 distinct experiments (Poitiers CHU) and the mean and SE of the 3 Ct were calculated.

Although the experiments were run over 3 days (at Poitiers CHU), the sampling from a given capsule gave constant results. In all the experiments, techniques and targets, the results obtained in the different laboratories were reproducible among all the capsules, showing that the encapsulated control RNA are suitable for their use as internal quality controls. The Ct for variants were strictly equivalent. The highest variability came from the analysis kits and target genes as demonstrated above with the controls in solution.

### Compatibility with a RT-LAMP procedure

SARS-Cov-2 genome was successfully amplified using the RT-LAMP assay from both RNA 2 low and RNA 2 high capsules with a high sensitivity (100% positive samples at 25 copies per reaction for RNA2 low, 40 copies for RNA2 high and 50% positive samples at 8 copies for RNA 2 high). Annealing temperature peaks were obtained for both S and N regions for 90% of the positive samples with annealing temperatures ranging between 88.4-88.7 °C and 83.4-83.8 °C, for N and S regions, respectively. For 10% positive samples only one annealing temperature peak (S or N) was obtained. This last result was principally obtained for low copy numbers. This was already observed in other RT-LAMP experiments involving other types of samples and is probably due to differences in primers binding probabilities to their respective targets.

In the field, 1000 copies per reaction were taken so that the signal came out early, validating the run before or at the same time as positive samples came out.

The results are given Table 3.

**Table 3.** RT-LAMP results from CIRAD.

| RNAsheII<br>RNA 2 | Copies / reaction | Amplification<br>time (min) | Annealing temperature |  |
| --- | --- | --- | --- | --- |
|  |  |  | N gene | S gene |
| Low | 1000 | 10.00 | 88.5 | 83.50 |
| Low | 500 | 11.75 | 88.5 | 83.7 |
| Low | 500 | 11.00 | 88.5 | 83.7 |
| Low | 250 | 12.75 | 88.5 | 83.6 |
| Low | 250 | 13.50 | 88.6 | 83.7 |
| Low | 200 | 14.50 | 88.6 | 83.8 |
| Low | 200 | 14.50 | 88.4 | 83.8 |
| Low | 200 | 16.75 | 88.7 | 83.7 |
| Low | 100 | 13.00 | 88.6 | 83.8 |
| Low | 100 | 12.50 | 88.8 | 83.7 |
| Low | 100 | 13.00 | 88.5 | 83.7 |
| Low | 50 | 13.50 | 88.7 | 83.8 |
| Low | 50 | 17.00 | 88.7 | 83.8 |
| Low | 50 | 14.00 | 88.7 | 83.7 |
| Low | 25 | 16.75 | 88.3 | 83.8 |
| Low | 25 | 14.00 | 88.6 | 83.8 |
| Low | 25 | 13.50 | 88.4 | 83.6 |
| High | 1000 | 10.30 | 88.7 | 83.7 |
| High | 1000 | 10.15 | 88.7 | 83.7 |
| High | 1000 | 10.15 | 88.6 | 83.7 |
| High | 200 | 12.00 | 88.6 | 83.7 |
| High | 200 | 11.30 | 88.6 | 83.6 |
| High | 200 | 11.30 | 88.6 | 83.6 |
| High | 200 | 10.45 | 88.7 | 83.7 |
| High | 40 | 12.30 | 88.7 | 83.7 |
| High | 40 | 13.45 | - | 83.7 |
| High | 40 | 13.15 | - | 83.7 |
| High | 40 | 13.45 | 88.6 | 88.5 |
| High | 8 | 19.45 | - | 83.4 |
| High | 8 | 23.30 | 88.7 | - |
| High | 8 | - | - | - |
| High | 8 | - | - | - |

RNA 2 samples High and Low were rehydrated with 50 μL of water and diluted to have between 1000 and 8 copies per reaction and amplified using the Runcov RT-LAMP protocol of the CIRAD La Réunion. The amplification time and annealing temperatures are given.

### “Real life” use of the capsules as internal quality controls

For the monitoring of SARS-CoV 2 detection in the field by RT-LAMP, the RNA 2 variant (1000 copies) was used as a positive control for 60 RT-LAMP tests run on samples taken and analyzed in 4 different sites (A, B, C, D). This control was successfully amplified in each run (Fig 8).

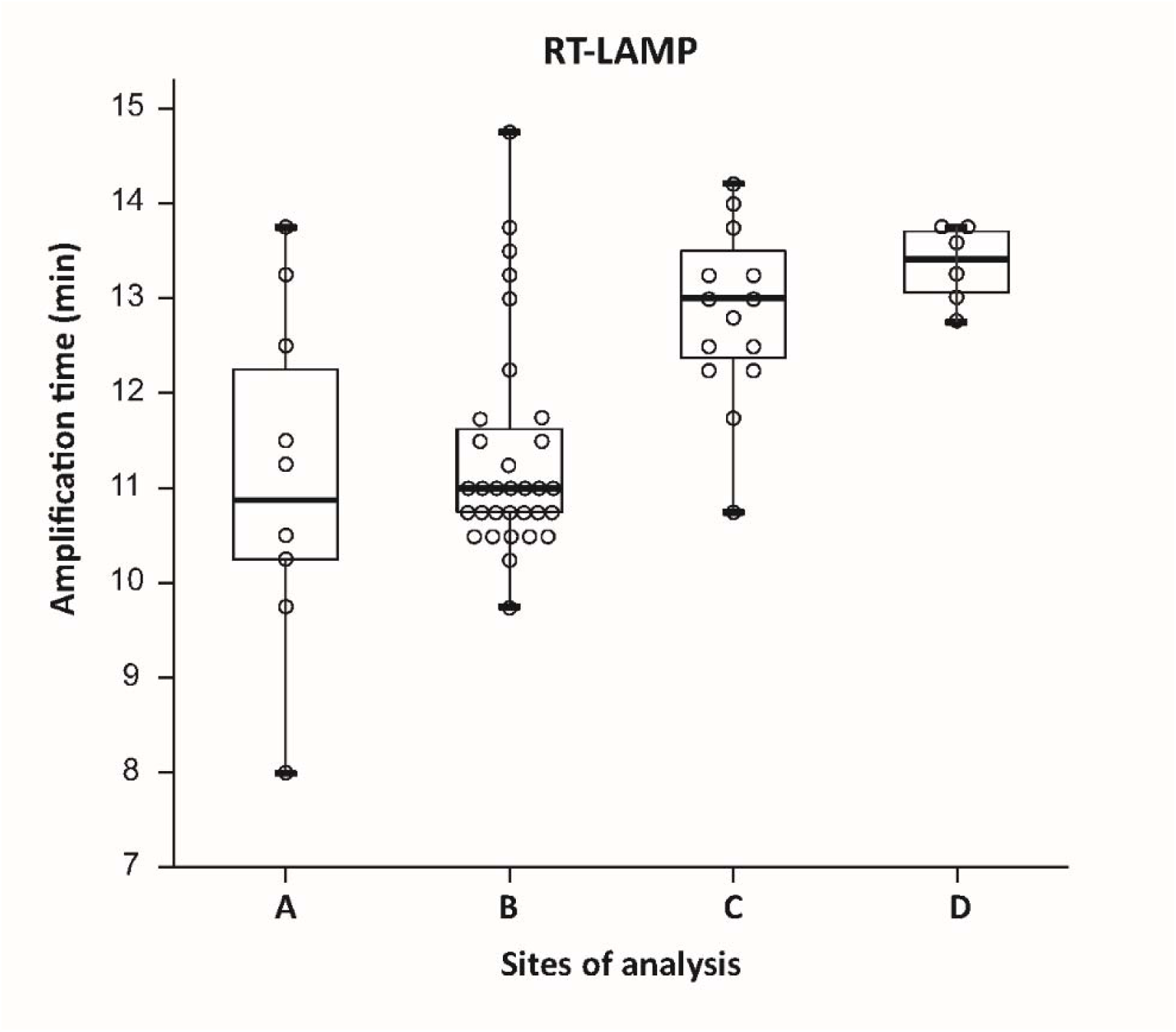

For each test, both N and S regions were amplified, generating annealing peaks within the expected annealing temperature ranges (not shown).

This field study shows that the capsules can be used as positive controls at the point of care or on site with simplified logistics due to their stability at room temperature.

A series of experiments was also conducted in a private clinical laboratory. The laboratory received the capsules and the instructions for the capsule opening and RNA rehydration. However, they were free to choose any volume above 10 µL. They included the rehydrated RNA in their routine diagnostic test. The results are shown Fig 9.

**Fig 9.**
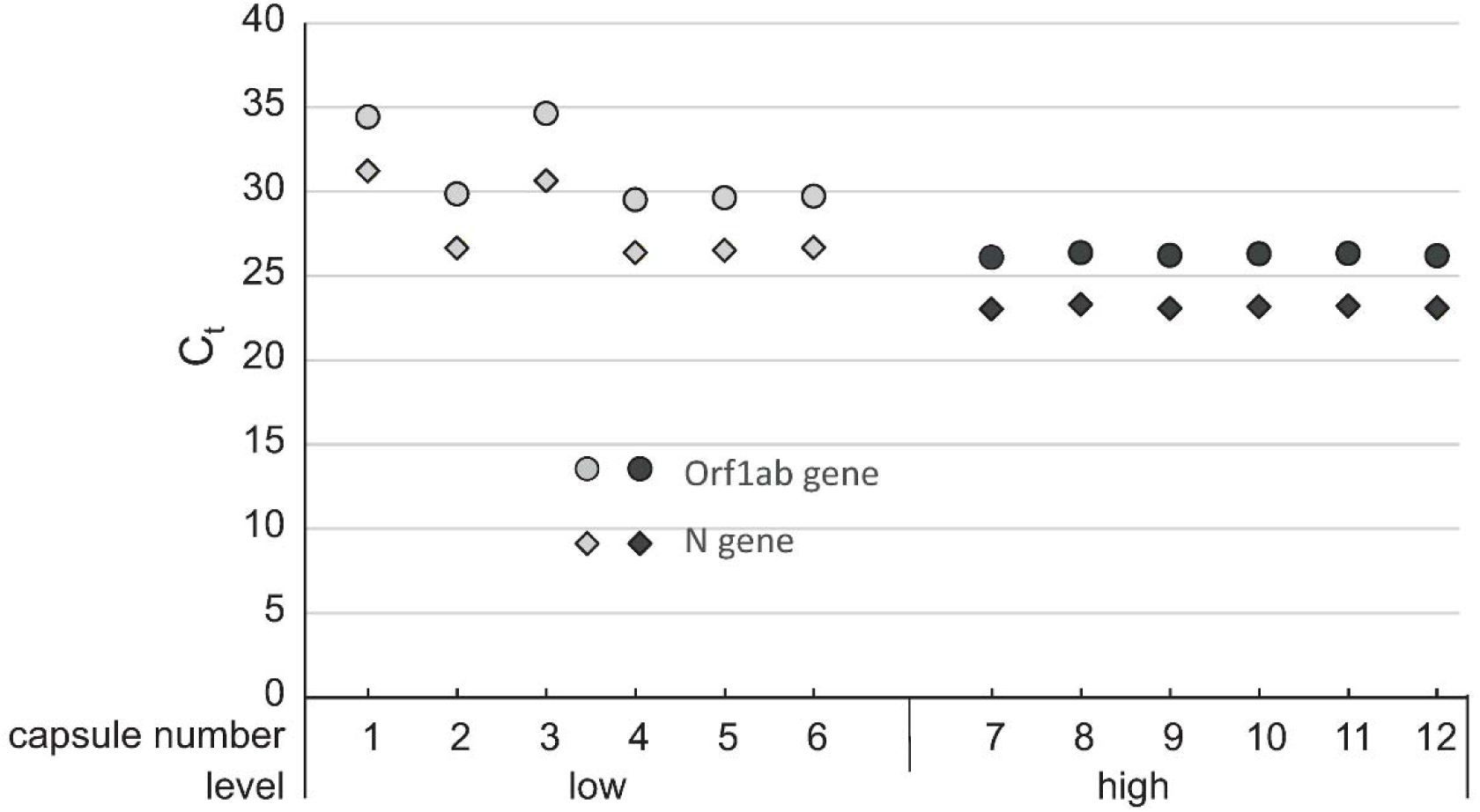
Private clinical laboratory. Results on 2 target genes and 2 levels on 6 capsules for each.

The results showed a good reproducibility among the capsules except for 2 capsules with a low RNA copy number. In the absence of a clear standard operation procedure like those usually provided in molecular diagnostic kits, the operator probably had rehydrated the RNA with a higher volume of water, decreasing the number of copies in the test.

So, the capsules can conveniently be used as reference materials for internal quality control of the monitoring of the diagnostics procedures, without the need for extensive training of the operators but with clear instructions to guarantee the reproducibility of the procedure for internal quality control purpose.

## Conclusion

Our aim was to demonstrate the suitability of encapsulation for the stabilization of positive control material for SARS-CoV-2 detection. We found that both 10,000 and 100,000 copies per capsule were sufficient for reliable RT-qPCR and RT-LAMP analysis. The signal was reproducible across different variants, targets, and analysis methods. Additionally, the RNAshell®-encapsulated RNA exhibited exceptional stability, with a shelf life of more than 10 years at room temperature. This stability presents several advantages: it simplifies the use of these controls in field settings, especially when temperature fluctuations are likely, and it enables the preparation and distribution of large, strategic batches that can be stored for extended periods and utilized for standardized testing procedures or across multiple detection sites.

## Data Availability Statement

All relevant data are within the paper and its Supporting Information file.

## Supporting Information

File containing all the data points obtained by the different laboratories using different amplification techniques and apparatuses on the 3 virus variants and their target genes.

## Supporting information

Raw Ct and LAMP data

## Acknowlegdements

We thank Twist bioscience for the gift of the RNA controls and suggestions for the writing of the paper.

